# Semantically Defined Subdomains of Functional Neuroimaging Literature and their Corresponding Brain Regions

**DOI:** 10.1101/157826

**Authors:** Fahd H Alhazmi, Derek Beaton, Hervé Abdi

**Affiliations:** School of Behavioral and Brain Sciences The University of Texas at Dallas MS: GR4.1 800 West Campbell Rd. Richardson, TX 75080 USA; Rotman Research Institute Baycrest Health Sciences 3560 Bathurst Street Toronto, ON, Canada M6A 2E1 School of Behavioral and Brain Sciences The University of Texas at Dallas MS: GR4.1 800 West Campbell Rd. Richardson, TX 75080 USA

**Author notes:** Authors note* The majority of the work was done while FA and DB were at The University of Texas at Dallas, and completed while DB was at the Rotman Research Institute. FA is now at The Graduate Center, CUNY. Correspondence can be addressed to any author. FA DB, HA.

**Keywords:** Meta-analysis, Neurosynth, Correspondence Analysis, Multivariate, Semantic Analysis, Recommendation engine

## Abstract

The functional neuroimaging literature has become increasingly complex and thus difficult to navigate. This complexity arises from the rate at which new studies are published and from the terminology that varies widely from study-to-study and even more so from discipline-to-discipline. One way to investigate and manage this problem is to build a “semantic space” that maps the different vocabulary used in functional neuroimaging literature. Such a semantic space will also help identify the primary research domains of neuroimaging and their most commonly reported brain regions. In this work, we analyzed the multivariate semantic structure of abstracts in *Neurosynth* and found that there are six primary domains of the functional neuroimaging literature each with their own preferred reported brain regions. Our analyses also highlight possible semantic sources of reported brain regions within and across domains because some research topics (e.g., memory disorders, substance use disorder) use heterogeneous terminology. Furthermore, we highlight the growth and decline of the primary domains over time. Finally, we note that our techniques and results form the basis of a “recommendation engine” that could help readers better navigate the neuroimaging literature.

## Introduction

Because terminology varies widely from study-to-study, and even more so from discipline-to-discipline, the neuroimaging literature is particularly difficult to synthesize. For example: (1) terminology changes over time (e.g., alcoholism to alcohol use disorders), (2) a single term can have many—or even amorphous—definitions (e.g., MVPA as multi-voxel or multivariate pattern analysis, which itself spans numerous different techniques), and (3) multiple terms describe the same concept (e.g., in vision studies “striate cortex,” “calcarine sulcus,” “V1,” “primary visual cortex,” and “Brodmann area 17,” all describe, essentially, the same brain region in functional neuroimaging). Such a diversity of terminologies makes interpretations, and even reviews, of the literature difficult to perform and consume.

To help navigate and consume results from the literature, several meta-analytic approaches (that link reported brain activations with keywords and topics) have been developed, such as coordinate-based meta-analysis (CBMA). CBMA was specifically developed for aggregating and synthesizing neuroimaging data reported in a standard format (P. T. Fox et al. 2014). Some of the most prominent CBMA tools used in research are BrainMap (Laird et al. 2005), SumsDB (Van Essen et al. 2009), Brede (Nielsen 2003) and NeuroSynth (Yarkoni et al. 2011)—the database of interest in this paper. The main functionalities of many CBMA tools are to (a) store coordinate information by study, and (b) provide spatially consistent meta-analytic activation maps. For example, Nielsen and colleagues (Nielsen et al. 2004) analyzed 121 neuroimaging papers with 2,655 reported activations loci using probability models followed by a non-negative matrix factorization-latent semantic analysis to associate brain coordinates with the words used in the papers’ abstract (e.g., “pain” was strongly associated with the anterior cingulate). More recently, Poldrack and colleagues (Poldrack et al. 2012) analyzed more than 5,800 papers to model the associations between topics derived from the full text of studies and the reported peak coordinates via “topic mapping.” Poldrack and colleagues showed that—with a “topic mapping” approach to semantic analysis—CBMA could reveal new relationships between brain activation and cognitive processes or psychiatric disorders (for different flavors of CBMA, see e.g., Rubin et al. 2016; de la Vega et al. 2016). Furthermore, some extensions of CBMA can link additional data types (e.g., gene expression) with brain regions and keywords (Mesmoudi et al. 2015; A. Fox et al. 2014; Rizzo et al. 2016).

Although the main functionalities of many of the CBMA tools are to (a) store coordinate information by study, and (b) provide meta-analytic activation maps (often based on terminology usage, e.g., which regions are most associated with “vision”, “anger”), CMBA tools fall short of revealing the primary domains of neuroimaging and the brain regions most associated with these domains (although CMBA tools are designed for that goal, i.e., meta-analyses). This limitation exists in part because of the absence of a common “semantic space” of the functional neuroimaging literature.

In this work, we define the primary domains of functional neuroimaging based on the semantics of the literature (i.e., abstracts)—a necessary step towards the definition of formal brain or cognitive ontologies. Our study is designed to achieve three major goals: (1) define a “semantic space” of the neuroimaging literature (which forms the basis of a “recommendation engine” to identify papers with high semantic similarity), (2) identify semantically defined domains within the literature, and (3) map brain activations onto these domains. To do so, we used correspondence analysis (CA)—a technique similar to principal components analysis (PCA) that was originally created for analyzing the co-occurrences of words in a corpus (Lebart et al. 1998; Benzécri 1976; Escofier-Cordier 1965, Abdi & Béra, 2014)—to identify neuroimaging domains from co-occurrences of words in the neuroimaging literature as identified in the Neurosynth database (Yarkoni et al. 2011).

First, we applied CA to a 10,898 studies × 3,114 words matrix; because CA on this matrix generates thousands of components, we used split-half resampling (SHR) to identify the most reliable and replicable components. Next, we applied hierarchical clustering (HC) within the subspace (i.e., the subset of components) identified by SHR to identify the primary subdomains in functional neuroimaging. We then investigated how these clusters change over time. Next, we generated brain maps (in MNI space) conditional to both the components and clusters, which highlight the brain regions most commonly associated with the components and clusters we identified. We also include a comparison of our brain maps with recent maps from Yeo et al., (2015). Finally, our work provides the basis of a “recommendation engine” that allows researchers to find semantically similar papers (based off of PubMed IDs).

## Methods

### Data and preprocessing

Neurosynth is an open source and open science initiative—hosted via the website www.neurosynth.org—that facilitates meta-analyses and reviews of the functional neuroimaging literature (Yarkoni et al. 2011). Neurosynth, at the time of this writing, contains more than 11,406 articles from the functional neuroimaging literature. When we began this work, Neurosynth contained 10,903 articles (from 43 journals), which were the basis of this study. As an aside, some articles in our data set no longer appear in Neurosynth because Neurosynth periodically updates its content for exclusion (e.g., to remove structural only studies) and public release. See http://github.com/neurosynth/neurosynth-data and http://www.neurosynth.org/ for details

Neurosynth uses automated webcrawlers to fetch data (e.g., abstract text, peak coordinates) and metadata (e.g., PubMed ID, title, year published, journal) of neuroimaging studies. For our study, we created and analyzed two data tables derived from Neurosynth data: (1) a “studies × words” matrix and (2) a “studies × voxels” matrix, where studies are identified by their PubMed ID (PMID) number. To achieve our three goals, our study comprised three major parts that correspond to each goal, wherein each major part has several steps. All analyses were conducted with a variety of publicly available packages (noted in relevant sections) or in-house scripts written in MATLAB (MathWorks, Natick, MA), R (R Core Team 2017), and Python (Python Software Foundation, https://www.python.org/) languages and environments.

To create a studies-by-words matrix for analysis, we acquired information from Neurosynth and PubMed (http://www.ncbi.nlm.nih.gov/pubmed/). With PMIDs from the Neurosynth database, we obtained from PubMed the text in all abstracts associated with the studies in the Neurosynth database. Next, we used the tm package in R (Feinerer 2011) to conduct several preprocessing steps that were used in previous works (Ailem et al. 2016) and that consisted in: (1) converting all text to lower-case, (2) removing all punctuations, numbers, emails and web addresses, (3) removing all words of length one or two, (4) removing step words, meta-words and words that describe numbers, quantities, nationalities, cities, or names (e.g., “publisher,” “article,” “date,” “ten,” “zero,” “weeks,” “european,” “montreal,” “welcome”), (5) converting British English to American English, (e.g., “behaviour” to “behavior”) and finally (6) stripping out white spaces. Once the data were cleaned, some words with different meanings were updated so they did not have the same “stems.” For example, “posit,” “positive,” “positively,” “position,” and “positioning” would correspond to the same stem “posit;” therefore some words were altered so they would only have the same stem if they had (in general) the same meaning. In a final step, we went through all remaining words individually to identify words that were potentially missed in the previous steps (e.g., “science,” “academic,” “publishing”). With a final set of words, we created a matrix of studies (rows) by words (columns). Each cell of this matrix contained the number of occurrences of a word used in the abstract of a study; for example, the abstract of PMID 17360197 used the word “cold” 28 times. Finally, we eliminated infrequent words in the studies-by-words matrix: words with frequencies below the 3^rd^ quartile (in our case: less than 16 occurrences) were removed. This step was followed by the removal of two studies that were withdrawn by the publisher. The final studies-by-words matrix contained 10,898 studies and 3,114 words (the full data table is provided in https://github.com/fahd09/neurosynth_semantic_map).

### Correspondence Analysis

Because our data are the number of occurrences (i.e., counts), we decided to use correspondence analysis (CA)—a technique designed specifically to analyze co-occurrences and often described as a version of PCA tailored for qualitative data. Like PCA, CA decomposes a matrix into orthogonal components rank-ordered by the variance they explained (Greenacre 1984; Abdi & Béra 2014; Lebart et al. 1984). CA is a bi-factor analysis that accounts for the relationships between and within the rows and the columns of a (contingency table) data matrix. CA assigns to each row (study) and column (word) item a “component score” (a.k.a. factor score) that reflects the amount of variance this item contributes to a given component. CA places emphasis on rare items so that they contribute a high amount of variance, while frequent items contribute little variance (see Greenacre 2017); this is particularly useful for our study because frequent words (e.g., “brain”) will be near the origin (i.e., zero) of the components whereas rare words (e.g., “polymorphism”) will be far from the origin and thus are the sources of variance for the components. CA is closely linked to the independence assumptions of χ^2^, which is proportional to the total variance decomposed by CA and therefore CA decomposes in orthogonal factors the pattern of deviation to independence of the data. Finally, because both rows and columns are represented in the same space (with the same variance), we can interpret the relationships within row items and within column items as well as the relative relationships between row and column items. Finally, because we wanted to identify brain regions most associated with semantically-defined domains, we used a technique called *supplementary projection* (also called “out of sample projection,” Greenacre 2017; Abdi & Béra 2014) that allows to predict a *supplementary* (i.e., new, or excluded) data set (i.e., studies × voxels) from the component structure of the *active* data set (i.e., studies × words).

We used in-house MATLAB code, as well as the ExPosition (Beaton et al. 2014) and ggplot2 (Wickham 2009) packages in R to perform CA as well as visualizations and additional analyses (i.e., visualizations, resampling-based inference tests, clustering, and supplementary projections; see following sections).

### Split-half Resampling

Split-half resampling (SHR, Strother et al. 2002; Churchill et al. 2012) is a cross-validation (CV) technique that evaluates the stability of the results of a statistical analysis performed on a data set by randomly splitting this data set into two (approximately) equal sized non overlapping data sets, and then performing the same analysis on each data set. The similarity (e.g., correlation) between the results obtained from these two data sets is then used to evaluate the reliability of the results (i.e., replicable effects). SHR is performed many times to create a distribution of reliability estimates.

We used SHR to identify the most replicable components in two ways: (1) split the data by study (rows), and (2) split the data by words (columns); in both approaches, we performed CA on each split set, and then computed the absolute correlation^1^ between the component scores of each split. SHR was performed 1,000 times to create a distribution of (absolute) correlations between components for both the (1) row component scores conditional to the columns, and (2) the column sets of scores conditional to the rows. We then computed the average (absolute) correlations to detect which components (after 1,000 splits) were most replicable between splits in order to identify a low rank approximation of the semantic space (i.e., component selection via SHR).

### Clustering of Studies and Assignment of Words

We performed hierarchical clustering (HC), with squared Ward linkage (Murtagh & Legendre, 2014), on the subset of reliable (as identified by SHR) component scores for the studies (rows). We chose squared Ward linkage because its objective function minimizes the error sums of squares (and thus provides an optimal ANOVA-like configuration). The component scores take into account the explained variance per component (i.e., Component 1 explains more variance than Component 2). After HC, we performed cluster stability analysis via Calinski-Harabasz index (Calinski & Harabasz, 1974) in order to identify a stable number of clusters. After the studies had been divided into clusters, we used distance-based classification in order to assign each word (column) to the closest study cluster barycenter (i.e., the point that represents the multidimensional mean of all studies in a given cluster). Hierarchal clustering and cluster stability were conducted in R via hclust and clusterCrit (Desgraupes 2015), respectively.

### Producing Brain Maps

Activation maps are represented in Neurosynth as peak activations of individual studies as centers of a sphere with a radius of 6mm. Voxels inside the sphere have a value of 1 and the other voxels have a value of 0. The voxels-by-studies matrix then uses a vectorized (flattened) version of the peak activation maps with reference to a 3D brain. The voxels-by-studies matrix initially contained 10,898 studies and 228,453 voxels (i.e., the voxels within MNI space). For our analyses, infrequently reported voxels (i.e., voxels that are reported in less than 10% of studies) were removed. The final studies-by-voxels matrix contained 10,898 studies and 206,077 voxels. We computed two different brain activation maps from the semantic space. The first type of activation map was a component-wise map. Brains were projected onto (i.e., predicted by) the semantic space—per replicable component—via supplementary projections. The second type of activation map was simply the sum of peak activations *per study cluster*.

### Supplementary Projections

Supplementary—a.k.a. out of sample—observations (or variables) can be integrated into an existing analysis performed on a different set of observations (or variables) referred to as the active data set. Supplementary data are assigned component scores by computing the least square projection for observations (or variables) onto the space defined by the active observations (or variables). We used supplementary projection to predict component scores for voxels *from* the component scores of studies defined in the in the semantic space (i.e., CA of studies × words). Predicted activation maps (from the supplementary scores) were projected back into MNI space. Functional volumes are then projected to the brain surface space using Caret5 software (Van Essen et al. 2001; http://brainvis.wustl.edu/) with the “Interpolated Voxel Algorithm.” All resulting maps (5 components maps and 6 cluster maps) are shared publicly in a Neurovault (Gorgolewski et al. 2016) repository: http://neurovault.org/collections/2002/.

## Results

CA was applied to a 10,898 studies × 3,114 words matrix and produced 3,112 components (see Figure 1a for the Scree plot). Split-half resampling (SHR) and cluster stability analysis revealed 5 reliable components (Figure 1b and c) and 6 reliable study clusters that define the latent semantic space of functional neuroimaging literature (via *Neurosynth*).

**Figure 1:**
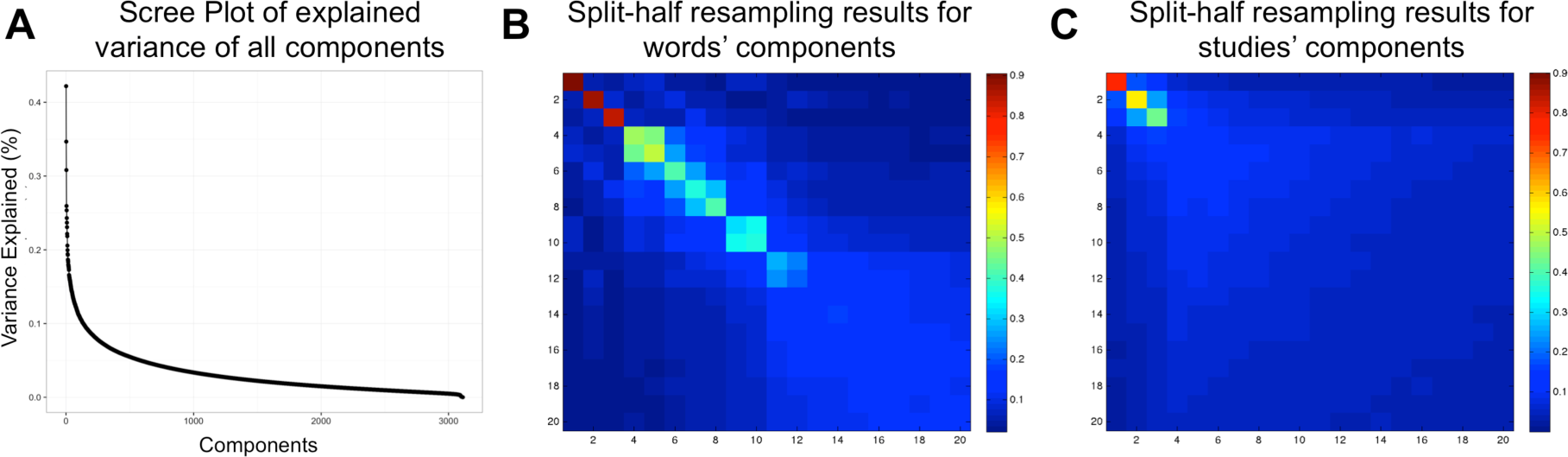
Variance explained and reproducibility (via split-half resampling; SHR) of latent semantic components. (a) The Scree plot shows the explained variance per component for all 3,112 components. Figures 1 (b) and (c) show heatmaps of correlations between component scores after SHR where (b) shows average (absolute) correlations after SHR for the words component scores and (c) shows the average (absolute) correlations after SHR for the studies component scores; only components 1 through 20 are shown. Both the Scree plot and the heatmap for the studies component scores suggest three high variance and highly reproducible components. The heatmap for the words component scores also show that the first three components are highly reproducible, but also that Components 4 and 5 are reproducible in the words component scores.

To help interpret the components of our semantic space, we used the words and studies at the extremes (i.e., highest contributing variance) for each component (Figure 3 for extreme words; Supplemental tables 1–5 for extreme studies). Table 1 shows the total and relative number of studies and words per cluster. As with the components, the most (and least) frequent words within each cluster help us interpret the cluster’s meaning (Supplemental tables 6). Furthermore, we also identified the words closest to the barycenter of each cluster (across all five dimensions; Supplemental table 7). We also provide the titles and PubMed IDs of the twenty studies closest to the barycenter of each cluster in Supplemental tables 8-13 as well as the overall “most average” and “most unique” studies and terms in Supplemental table 14. Component maps—which present two components at a time—are presented in Figure 2. We present component maps of the words and studies separately. In each map, we color each dot (i.e., a study or word) by its associated cluster. Components 1, 2, and 3 are visualized in Figures 2a–d. We show Components 4 and 5 separately from the other components (Figures 2e and f) because studies on Components 4 and 5 constitute a single cluster (see next section). Brain maps for the components are presented in Figure 3, and brain maps for clusters are presented in Figure 4. In *Results,* the components and clusters are first referred to by numbers: The component number reflects its rank order (by variance), but the cluster numbers are arbitrary. We provide interpretations of components and names for clusters after their descriptions.

**Figure 2:**
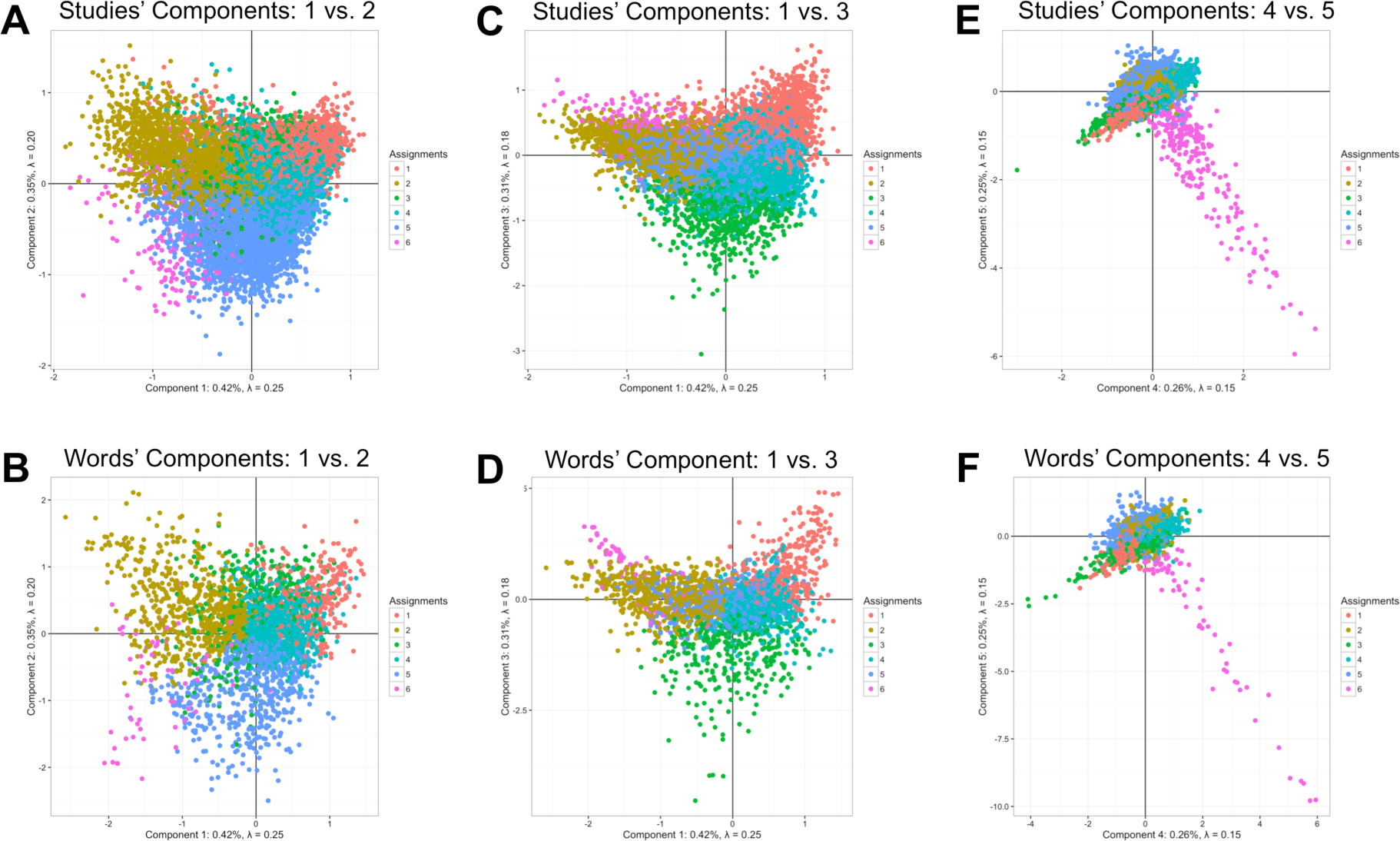
Visualization of the semantic space of functional neuroimaging literature. Component scores for both the words and the studies on Components 1 through 5 are visualized in a series of 2D figures. Axes are components and individual dots represent either a particular study or particular word. Words and studies are colored by which cluster they belong to and thus illustrate the large sub-domains within *f*MRI. Figures 2 (a) and (b) show the words and studies (respectively) component scores for Components 1 (horizontal) and 2 (vertical). Figures 2 (c) and (d) show the words and studies (respectively) component scores for Components 1 (horizontal) and 3 (vertical). Figures 2 (e) and (f) show the words and studies (respectively) component scores for Components 4 (horizontal) and 5 (vertical). While most words and studies form large groups within the axes, Components 4 and 5 show a highly specific subset of words and studies, of which nearly all are assigned to Cluster 6 (typically fMRI studies that include genetic and molecular terms).

**Table I:** The total number of studies and terms per cluster. *Note.* Our analyses revealed six clusters. The total number of studies and terms per cluster are provided. Furthermore, we provide a description that helps characterize the contents of each cluster.

| Cluster | # Studies (%) | # Terms (%) | Brief description |
| --- | --- | --- | --- |
| 1 | 1,569 (14.4%) | 351 (11.3%) | Knowledge Representation & Language Processing |
| 2 | 2,272 (20.8%) | 720 (23.1%) | Development, Lifespan and Disorders |
| 3 | 728 (6.7%) | 347 (11.1%) | Sensation, Movement and Action |
| 4 | 3,159 (28.9%) | 1019 (32.7%) | Cognition & Psychology |
| 5 | 2,927 (26.9%) | 612 (19.7%) | Decision, Emotion, & Substance Use |
| 6 | 243 (2.2%) | 65 (2.1%) | Imaging Genetics |
| *All* | *10,898* | *3,114* |  |

**Figure 3:**
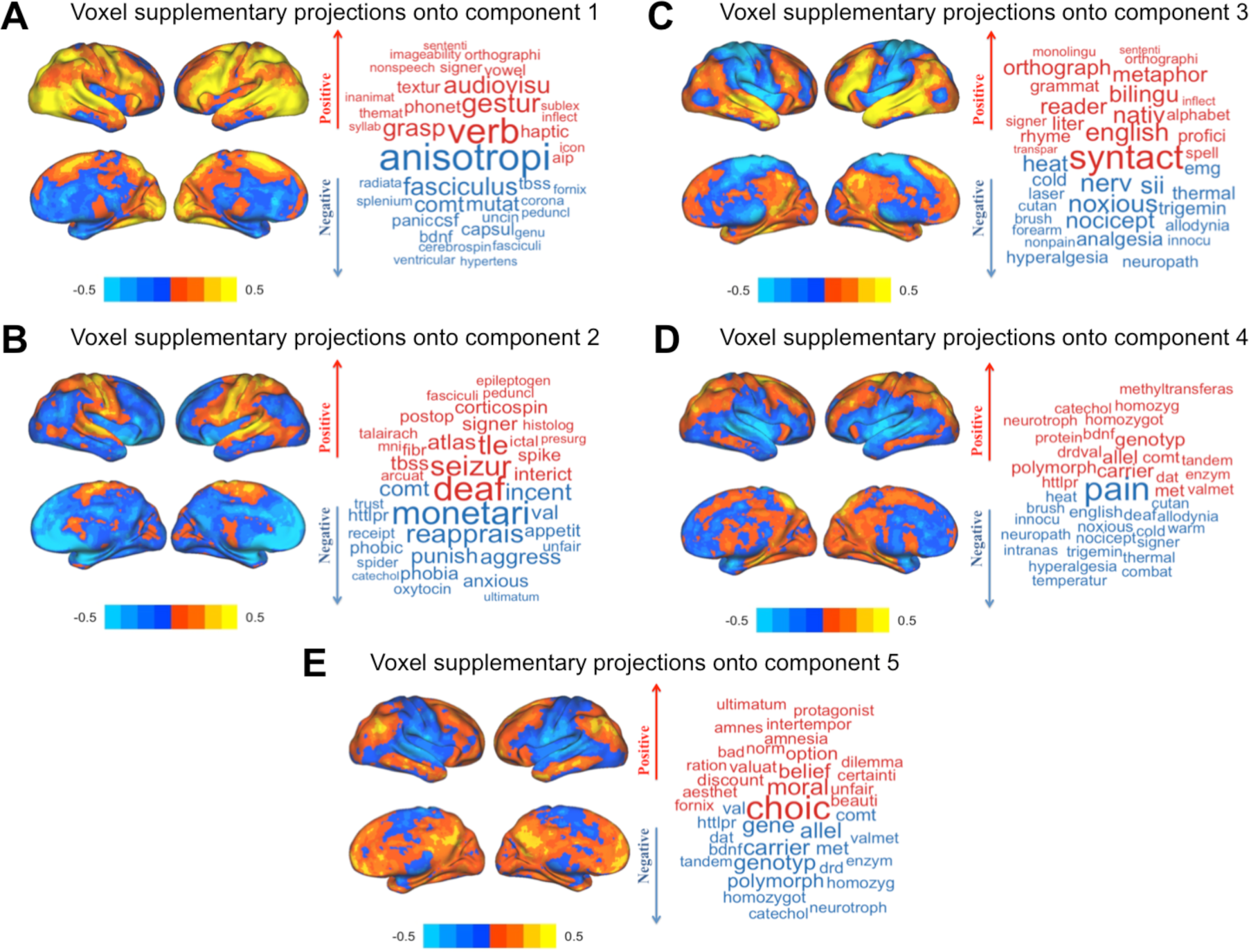
Visualization of brain maps, per component, as predicted (via supplementary projections) by the words × studies component scores (left) and a word cloud that shows some words that either loads on the positive or negative axis (right). (a) The projected map for Component 1 shows that the positive side (marked in red) is associated with the left temporal lobe, bilateral occipito-temporal, and parietal regions, while the negative side (marked in blue) is associated with many subcortical structures. (b) The projected maps for Component 2 show that the positive side is associated with bilateral somatosensory areas and the right cerebellum (not rendered in surface plots), while the negative side is associated with subcortical structures as well as medial prefrontal cortex. (c) The projected brain maps for Component 3 show that the positive side is associated with the left lateralized language-related areas while the negative side is associated with somatosensory cortex in addition to the brainstem (not rendered in surface plots). (d) The projected map for Component 4 shows that the positive side is associated with medial structures of parietal areas (precuneus) while the negative side is associated with bilateral somato-sensory areas as well as the insular cortex and brainstem (not rendered in surface plots). (e) The projected brain maps for Component 5 show that the positive side is associated with the posterior cingulate and medial prefrontal cortices while the negative side is associated with bilateral somato-sensory areas and the brainstem (not rendered in surface plots).

**Figure 4:**
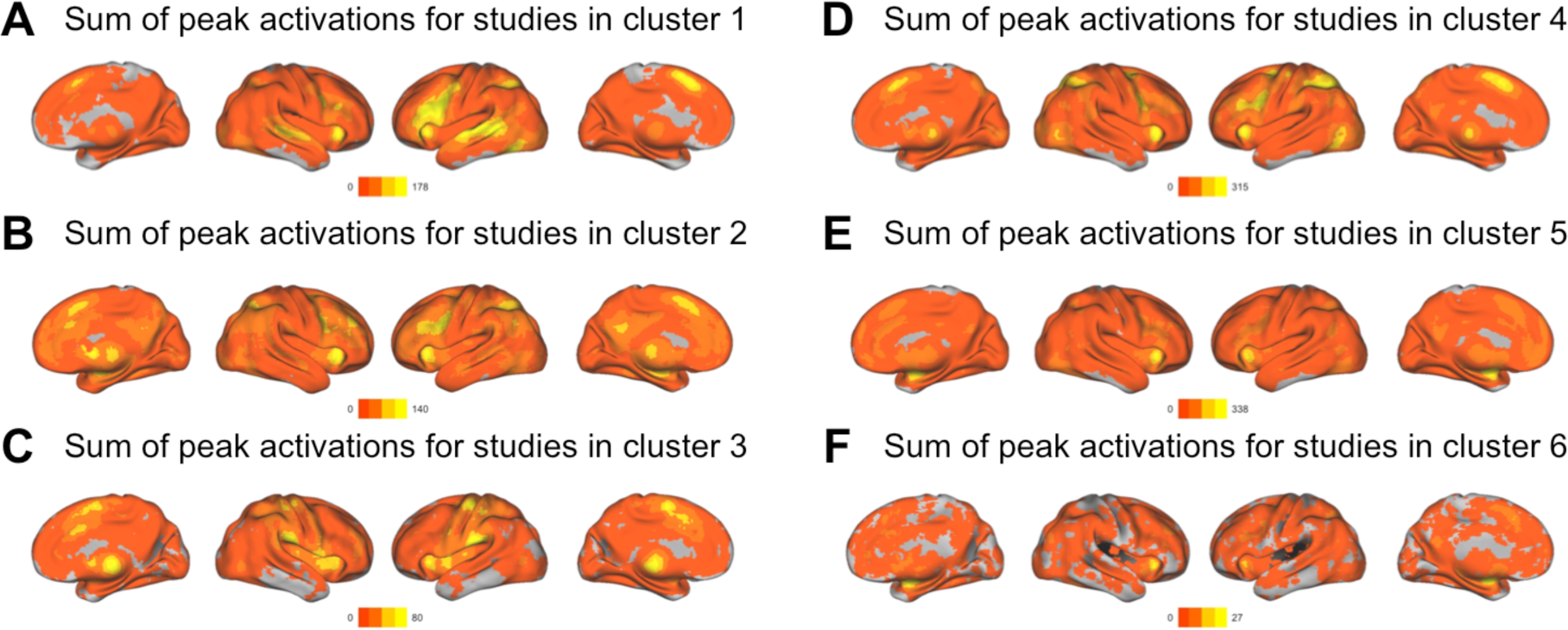
Visualization of brain maps, per cluster, computed as the sum of peak activations for all studies within a particular cluster. (a) Summed activations of individual studies in cluster 1 (*Knowledge Representation and Language Processing*) showed in the bilateral peaks in the frontal and temporal lobes—which are often associated with language and knowledge representation—in addition to a peak area in the anterior cingulate cortex. (b) Summed peak activations of studies in cluster 2 (*Development, Lifespan and Disorders*) showed a diffuse pattern of reported activations in frontal and parietal areas, as well as subcortical regions. (c) Summed peak activations of studies in cluster 3 (*Sensation, Movement and Action*) showed in bilateral somatosensory areas and the thalamus. (d) Summed peak activations of studies in cluster 4 (*Cognition and Psychology*) showed in the mid-line and bilateral areas in frontal regions, in addition to bilateral occipito-temporal regions. (e) Summed peak activations of cluster 5 (*Decision, Emotion, and Substance Use*) appeared in subcortical areas and medial frontal regions. (f) Summed peak activations of Cluster 6 (*Imaging Genetics*) showed almost entirely subcortical areas.

### Components

Component 1’s words and studies can be seen in Fig. 2a–b. Words at extreme positive and negative sides of Component 1, as well as the projected brain maps for Component 1 are shown in Fig. 3a. The projected brain map for Component 1 (Fig. 3a) show that: (1) The positive side of Component 1 is associated with the left temporal lobe, bilateral occipito-temporal, and parietal regions, and (2) the negative side of Component 1 is associated with many subcortical structures. Component 1 generally reflects basic science research on the positive side to clinical/translational neuroimaging research on the negative side (Figs 2a-b). While the basic science research is more associated with cortical structures, the clinical/translational research is more linked with subcortical structures (Fig 3a).

Component 2’s words and studies can be seen in Fig. 2a–b. Words at the extreme positive and negative sides of Component 2, as well as the projected brain map for Component 2 are shown in Fig. 3b. The projected brain maps for Component 2 show that: (1) the positive side of Component 2 is associated with bilateral somatosensory areas and the right cerebellum, and (2) the negative side of Component 2 is associated with subcortical structures as well as medial prefrontal cortex. Component 2 generally reflects a methodological spectrum that ranges from cognitive tasks on the negative side to multi-modal imaging (e.g., structural-functional association) on the positive side. The negative—and more densely populated—side of Component 2 is more associated with studies on affect and emotion and similar study types (Fig. 2a–b) with projections in subcortical and prefrontal cortex, whereas the positive side of Component 2 is linked to studies that rely on multi-modal imaging as well as other related methodologies and somatosensory, temporal and cerebellar projections (Fig. 3b).

Component 3’s words and studies can be seen in Fig. 2c–d. Words at the extreme positive and negative sides of Component 2, as well as the projected brain maps for Component 3 are shown in Fig. 3c. The projected brain maps for Component 3 show that: (1) the positive side of Component 3 is associated with the left lateralized language-related areas (e.g., temporal and frontal areas known as Broca’s Area and Wernicke’s Area), and (2) the negative side of Component 3 is associated with somatosensory cortex in addition to the brainstem. Component 3 generally reflects low-to-high level of cognition; studies about higher-order cognitive processes (especially linguistics) are on the positive side of Component 3 (with cortical projections to the bilateral temporal and frontal regions; Figs 2c–d; Fig. 3c), whereas studies on lower-order cognitive processes (such as sensation, perception, and direct sensory stimulation) are on the negative side of Component 3 (and also projects to the bilateral somatosensory areas and the brainstem in addition to the left cerebellum; Figs 2c–d; Fig. 3c).

Components 4’s and 5’s words and studies can be seen in Fig. 2e–f and show a highly distinct pattern from the other components, mostly driven by words and studies related to molecular, genetic, and genomic neuroimaging studies. Words at the extreme positive and negative sides of Components 4 and 5, as well as the projected brain maps for Components 4 and 5 are shown in Figs. 3d-e. The brain maps for Component 4 (Fig. 3d) show that: (1) the positive side of Component 4 is associated with medial structures of parietal areas (precuneus), and (2) the negative side of Component 4 is associated with bilateral somato-sensory areas as well as the insular cortex and brainstem. The brain maps for Component 5 (in Fig. 3e) show that: (1) the positive side of Component 5 is associated with the posterior cingulate and medial prefrontal cortices, and (2) the negative side of Component 5 is associated with bilateral somato-sensory areas and the brainstem. Components 4 and 5 showed a distribution of words and studies that makes a near perfect 45° angle between Components 4 and 5 that extended out from the origin. These words and studies were almost entirely molecular, genetic, and genomic neuroimaging (i.e., “imaging genetics”) studies. While studies on Component 4 are more associated with genetic contributions to cognition in healthy populations, by contrast, the projections to parietal and frontal regions studies on Component 5 were more associated with genetic contributions in disordered populations with projections to bilateral somatosensory regions.

### Clusters

Cluster 1 contains studies/words primarily associated with language/speech production, comprehension, and disorders, as well as knowledge processing. Some examples include: *decod, word, superior, auditori, languag, semant, percept, speech, recognit, complex* (see Supplemental tables 6, 7, and 8). Figures. 2a-b show that this cluster primarily lies on positive sides of Components 1 and 3. Summed peak activations of individual studies in this cluster are localized in the bilateral frontal and temporal regions—which are often associated with language and knowledge representation—in addition to the anterior cingulate cortex (see Fig 4a). Cluster 1 represents studies that mainly investigate knowledge representation and language processing and we henceforth refer to Cluster 1 as *Knowledge Representation and Language Processing*.

Cluster 2 contains studies/words associated with developmental, lifespan, and aging studies, as well as their respective disorders. Some examples include: *patient, differ, chang, healthi, age, structur, breakdown, degen, ecnp, epidemiolog* (see also Supplemental tables 6, 7, and 9). Figs. 2a–b shows that this cluster primarily loads on the negative side of Component 1 and on the positive side of Component 2. Summed peak activations of studies in this cluster (see Fig 4b) showed a diffuse pattern of activations in the frontal and parietal areas, as well as in the subcortical regions. Cluster 2 represents studies that mainly investigate developmental and adult lifespan research in addition to brain disorders and we henceforth refer to Cluster 2 as *Developmental, Lifespan, and Disorders*.

Cluster 3 contains studies/words primarily associated with sensation (cutaneous and olfaction) and movement. Some examples include: *motor, pain, movement, hand, stimul, sensori, thalamus, somatosensori, reflex, anesthet* (see also Supplemental tables 6, 7, and 10). Figs. 2c-d shows that this cluster loads primarily on the negative side of Component 3. Summed peak activations of studies in this cluster (Fig. 4c) showed in the bilateral somato-sensory areas and the thalamus. Cluster 3 represents studies that mainly investigate sensation, movement and action and we henceforth refer to Cluster 3 as *Sensation, Movement, and Action*.

Cluster 4 contains studies/words associated with more “traditional” aspects of human cognitive neuroscience: those rooted in cognitive and experimental psychology (i.e., they rely primarily on behavioral tasks to examine neural correlates). Some examples include: *activ, function, task, area, fmri, network, memori, effect, visual, decay* (see also Supplemental tables 6, 7, and 11). Figures 2a–c show that Cluster 4 is closest to the origin point across all components with no apparent trend toward any axis. Summed peak activations of studies in this cluster (shown in Fig. 4d) showed in the mid-line and bilateral frontal regions, in addition to bilateral occipito-temporal region. Cluster 4 represents the majority of cognition and psychological-based functional neuroimaging research and we henceforth refer to Cluster 4 as *Cognition and Psychology*.

Cluster 5 contains studies/words that describe affective processes, such as emotional responses and decision-making, but also includes a number of studies and words related to substance use disorders and mood disorders. Cluster 5 includes the words: *emot, prefront, reson, cingul, examin, medial, amygdala, negat, social, diminish, take* (see also Supplemental tables 6, 7, and 12). Figs. 2a–b show that this cluster lies mostly on the negative side of Component 2. Summed peak activations of Cluster 5 (Fig. 4e) appeared in subcortical areas and medial frontal regions. Cluster 5 represents studies that mainly investigate decision-making, emotions and substance use (or abuse) and we henceforth refer to Cluster 5 as *Decision, Emotion, and Substance Use*.

Cluster 6 loads almost entirely and exclusively on both Components 4 and 5 (see Figs 2e–f). Cluster 6 contains words such as: *variat, genet, dopamin, gene, carrier, allel, genotyp, receptor, polymorph, dopaminerg, comt, serotonin, apo, norepinephrine* (see also Supplemental tables 6, 7, and 13). Summed peak activations of Cluster 6 (Fig. 4f) showed almost entirely in the subcortical areas. Cluster 6 represents a unique dimension (i.e., Components 4 and 5) of molecular, genetic, and genomic neuroimaging (“imaging genetics”) studies and we henceforth refer to Cluster 6 as *“Imaging Genetics”*.

### Temporal effects of clusters

Upon completion of the analyses, there were two clusters that stood out: (1) Cluster 4 (*Cognition and Psychology*)—which is essentially the “average” neuroimaging study because it is centered roughly on the origin of the components—and (2) Cluster 6 (*“Imaging Genetics”*)—which is comprised of the studies that define Components 4 and 5). Notably, Cluster 4 (*Cognition and Psychology*) reflects the origins of neuroimaging use (i.e., cognitive psychology), whereas Cluster 6 (*“Imaging Genetics”*) reflects the current state-of-the-art (i.e., translational and interdisciplinary work).

Figure 5 shows the relative frequency of the number of studies in each cluster sorted by year. Cluster 4 (*Cognition and Psychology*) accounts for a substantial amount of studies in the earlier years. For example, in the year 2000, approximately 50% of all neuroimaging studies (in Neurosynth) were in Cluster 4 (*Cognition and Psychology*). On the other hand, Cluster 5 (*Decision, Emotion, & Substance Use*) started as a small proportion of all neuroimaging studies in the earlier years, but now accounts for nearly 33% of all studies. We discuss the temporal properties of these clusters further in the Discussion.

**Figure 5:**
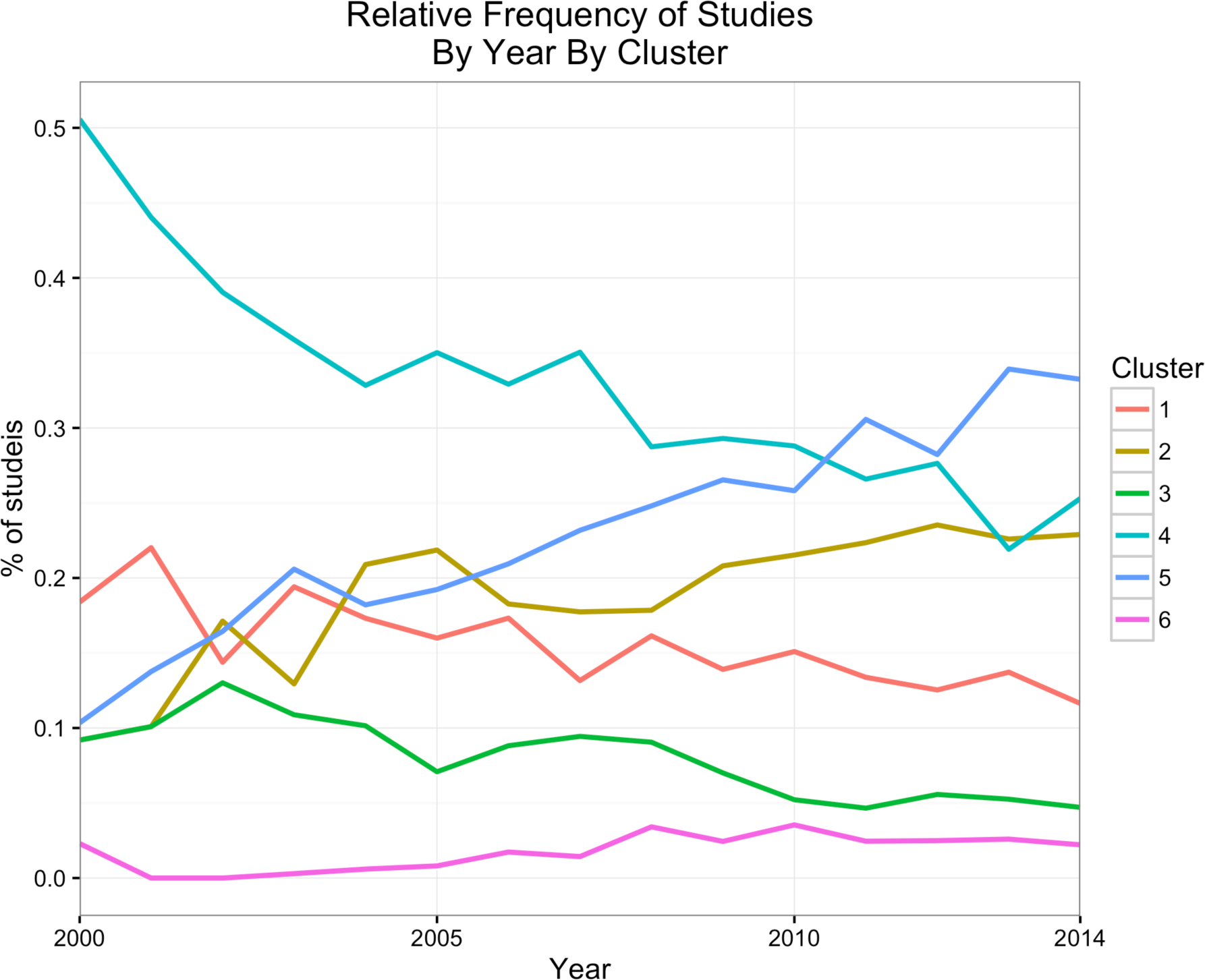
The proportion of studies within each cluster over time. Cluster 4 (*Cognition and Psychology*) was, and still generally is the core of *f*MRI research and as such comprises a substantial proportion of the literature. Though Cluster 4 (*Cognition and Psychology*) remains very large, it has decreased over time. Both Cluster 5 (*Decision, Emotion, Substance Use*) and Cluster 2 (*Developmental, Lifespan, Disorders*) have shown a considerable increase over time and now comprise, respectively, comparable proportions of the literature as Cluster 4 (*Cognition and Psychology*).

### Correlations with maps in Yeo et al. (2015)

In Yeo et al. (2015), a hierarchal Bayesian model was applied to 10,449 experimental contrasts in the BrainMap database in order to estimate the probability that each pre-defined task category would engage a specific cognitive component, and the probability that each cognitive component would engage brain regions (represented by voxels). Correlations between our component and cluster maps and Yeo et al. (2015)’s 12-component cognitive maps were computed using a custom script. We first downloaded the maps from *Neurovault* (http://neurovault.org/collections/866/, last accessed June 7, 2017). We only included the non-zero voxels from the component maps to exclude all non-valid voxels (i.e., outside the brain).

Figure 6 shows the correlations between our maps and the maps from Yeo et al (2015). We refer to Yeo et al.’s (2015) components as, e.g., Yeo Component 1 (YC1) or Yeo Component 6 (YC6) while we refer to our own components as “Component 1” or “Component 6.” There were several correlations of note for both the components (Figure 6a) and the clusters (Figure 6b). To note, although the magnitudes of those correlations are interpretable, the sign (or direction) of the correlation are not easily interpretable.

**Figure 6:**
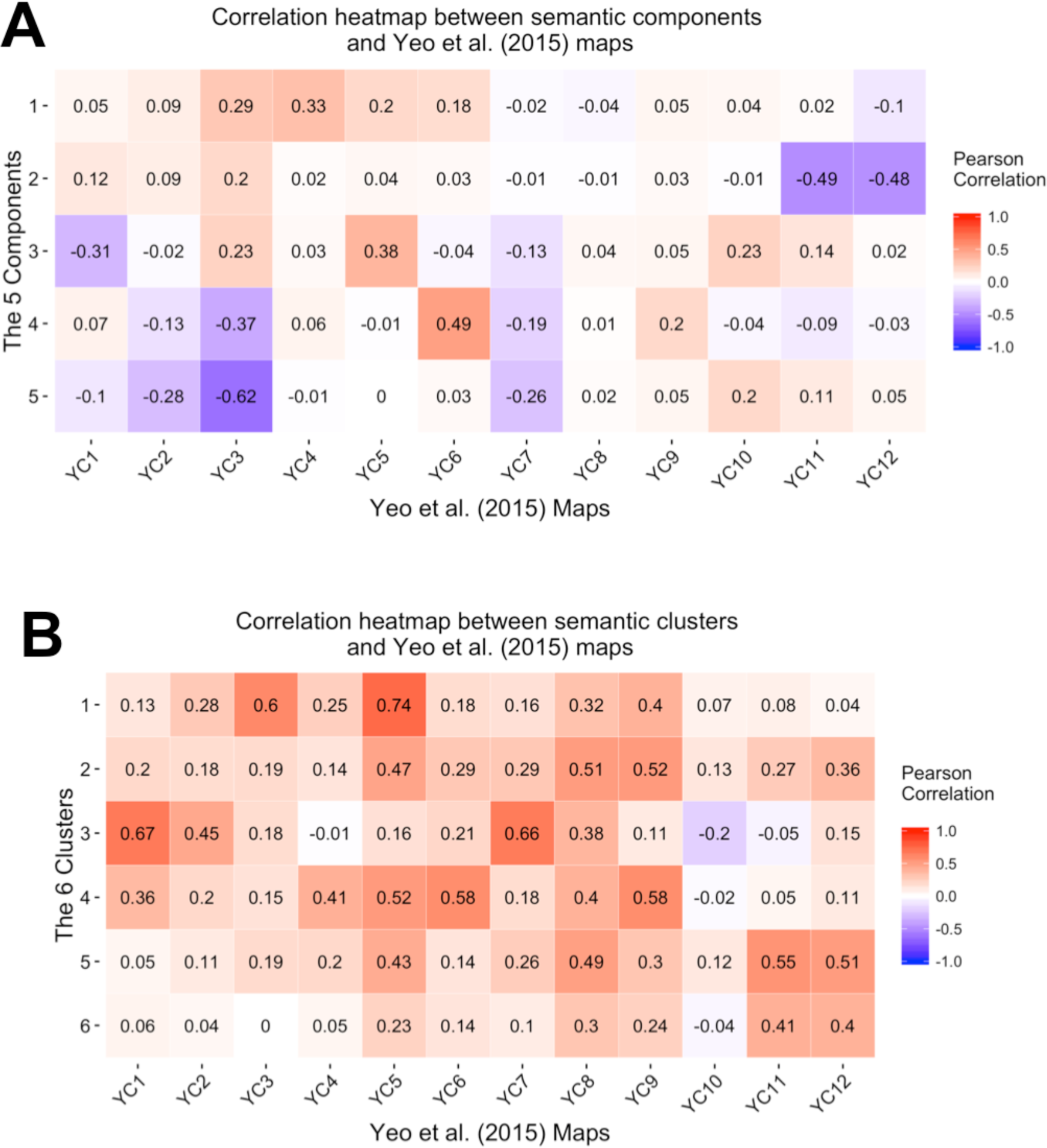
Shows the correlations between the maps generated by Yeo et al., (2015) and (a) our components or (b) our clusters. There are some notably high similarities between our brain maps (which were generated conditional to the latent semantic space) and the Yeo et al., (2015) maps, such as the Yeo components 11 and 12 with our Component 2 (see a), and the Yeo components 1 and 7 with our Cluster 3 (see b).

Figure 6a shows strong correlations between our Component 2 and YC11 and YC12. Both maps show strong association with bilateral subcortical structures (e.g., amygdala and striatum) in addition to their relationship with subcortical-related functions such as emotions and affect. Also, there is a strong correlation between our Component 3 and YC5 because both maps show strong association with temporal and frontal activations in addition to their relationships with semantic knowledge and language processing. Furthermore, there is a strong correlation between our Component 4 and YC6 because both maps show strong associations with medial parietal and frontal areas (commonly known as the frontal-parietal network; Smith et al. 2009).

Similar to the components, correlations between our clusters and the Yeo components are illustrated in Figure 6b. Our Cluster 1 is most correlated with YC 5 followed by YC3, all of which have activations in the temporal and frontal regions and are generally involved with knowledge representation and higher-order semantic processing. Our Cluster 3 is correlated with YC1 followed by YC7, both of which have activations in somatosensory areas and are involved in sensation and movement processing. Our Clusters 5 are both correlated with YC11 and YC12, all of which are associated with activations in subcortical structures and are associated with tasks that involve some aspect of affective or emotional processing. Although our Cluster 6 also share the same regions (and is most similar to YC11 and YC12), it comes mostly from molecular and genetic studies.

### Recommendation Engine

Finally, we provide a simple tool in R as a Shinyapp that works as a “recommendation engine” akin to preference and ratings systems (e.g., for movie preferences, shopping, or internet searches). While the app has many current and planned features, we only discuss the recommendation portion here. Our recommendation tool uses a distance-based search to retrieve papers (PMIDs) that are the most semantically similar to a given paper (PMID). Specifically, users only need to provide: (1) a PMID of a paper of interest, and (2) how many *N* similar papers to select. Our recommendation tool then provides the *N* papers closest to the target paper. Additionally, in the same way as for the papers, users can input a term of interest and retrieve the *N* closest terms. In both cases we also provide additional information such as to which primary domain (cluster) a study or term belongs. The recommendation is based on the component scores of the first five components (see Results). This recommendation engine, however, works based on the results in this paper and not the most recent version of *Neurosynth*. However, we provide code for all of our work including the recommendation tool and thus all results and recommendations can be updated as *Neurosynth* is updated. The recommendation engine with a brief description and how-to are available at the following address: http://bit.ly/neurosanity.

## Discussion

In recent years, there have been many meta-analyses, mega-analyses (analyses of pooled data across many studies), and other large-scale analyses of data within neuroimaging. In general, the aims of such analyses are to (1) test or refute findings and hypotheses (Wager et al. 2007), (2) build a consensus around particular models, hypotheses, or theories (Salimi-Khorshidi et al. 2009), (3) estimate consistency of findings (Wager et al. 2009), (4) help define related brain regions and networks (Toro et al. 2008; Mesmoudi et al. 2013), (5) interpret functional maps (Laird et al. 2011), or (6) segment the brain in new ways with resting-state *f*MRI measurements (Yeo et al. 2011; Power et al. 2011) or using massive multi-modal data (Glasser et al. 2016).

Using the meta-analytic cognitive component maps from Yeo et al. (2015) as a reference point to compare with our maps we showed a substantial overlap between many of our maps and maps from Yeo et al. (2015). However, our meta-analytic maps were predicted from the semantic space (i.e., abstracts) of the functional neuroimaging literature, whereas other authors took a more brain-centric approach, for examples: network- and meta-maps generated by with resting state *f*MRI (Yeo et al. 2011; Power et al. 2011) or via meta-analysis of data from hundreds or even thousands of studies (Yang et al. 2016; Yeo et al. 2015; Poldrack et al. 2012; de la Vega et al. 2016).

Our components explain the primary sources of variance of language used: in the field at large (i.e., Component 1), for methodological tools (i.e., Component 2), in various aspects of cognition (i.e., Component 3), and in relatively new studies with highly-specific terminology (i.e., Components 4 and 5). With supplementary projections we also showed that these language-based components are frequently associated with particular reported brain regions. While the components indicate language variation and gradients, our clusters define the boundaries of functional neuroimaging into specific—albeit large—sub-domains. Furthermore, our analyses revealed that there are, perhaps, biases or preferentially studied brain areas per domain (i.e., clusters).

Parts our semantic space also reflect, to a degree, current debates such as the distinctions between “neurological vs. psychiatric brain disorders” (White et al. 2012). For example, Crossley et al. (2015) recently used CBMA of voxel-based morphometric (VBM) studies to show a “neuroimaging-based” evidence for the biological distinctions between neurological vs. psychiatric disorders (Crossley et al. 2015). Our components show that neuroimaging studies in neurology and psychiatry do not use the same terminology and thus could be a source of the “versus” argument between neurological and psychiatric studies with respect to reported brain regions. As an illustration of this contention, we have selected some of the same neurology and psychiatry related terms used by Crossley et al., (2015) to highlight particular features of our components. First, all the words related to psychiatric or neurological disorders (Supplemental Table 15) appear on the negative side of Component 1—a configuration that supports our interpretation of a spectrum from basic science to applied and clinical neuroimaging. Furthermore, the neurological and psychiatric terms from Crossley et al., (2015) oppositely load on both Components 2 and 4 (Supplemental table 15): a configuration that reflects overall differences in *patterns* of terminology between neurological and psychiatric studies and thus expresses a dissociation of neurological studies and their regions (such as sensorimotor cortices and insula; in red) from psychiatric studies and their regions (such as limbic and prefrontal areas; in blue) as seen in Figure 3b. Further discrepant terminology can be seen in Supplemental Material (see Supplemental table 16).

Furthermore, the positions—and contents—of our clusters reveal a broad configuration of the neuroimaging literature. Cluster 4 (*Cognition and Psychology*) is the closest to the barycenter (origin of the axes across all components) and thus represents the average or most common neuroimaging study. This interpretation is supported by Cluster 4 (*Cognition and Psychology*) because it contains a substantial proportion of words and studies (~33% of words and ~29% of studies, see Table 1). Thus, much of the neuroimaging literature has been—and appears to still be—rooted in the approaches from cognitive and psychological domains. Summed peak activations of studies in this cluster (shown in Fig. 4d) show a high association with a wide set of cortical areas in the medial and bilateral frontal, occipital and subcortical regions that are associated with task performance. We also see opposition of clusters and this suggests that these are the sources of variance for our components. For example, Cluster 5 (*Decision, Emotion, and Substance Use*) is opposed to all other clusters on Component 2 (Figs. 2a-b)—a pattern that further supports the neurological vs. psychiatric dissociation of Component 2. Summed peak activations of studies in this cluster (shown in Fig. 4e) show high association with the subcortical areas and medial frontal regions that are generally associated with emotional processing and decision-making process. Similarly, Cluster 3 (*Sensation, Movemen,t and Action*) is opposed to all other clusters on Component 3 (Figs. 2c-d)—a component that, as we previously noted, expresses a spectrum from low-to-high level processing. Summed peak activations of studies in this cluster (shown in Fig. 4c) show high association with the bilateral somatosensory areas and the thalamus. Furthermore, Cluster 6 (*“Imaging Genetics”*) is almost entirely defined by the unique configuration of both Components 4 and 5 (Figs. 2e–f). Not only does Cluster 6 reflects a unique subfield of neuroimaging, but it also indicates that “imaging genetics” uses an almost exclusive set of words, different from the vocabulary of the rest of neuroimaging (cf., the 45 degree angle from Components 4 and 5). Summed peak activations of Cluster 6 (Fig. 4f) are almost entirely associated with subcortical areas. Finally, both Clusters 4 (*Cognition and Psychology*) and 5 (*Decision, Emotion, and Substance Use*) proportionally explain over half of the literature at any given time (Fig. 5).

Our clusters and their respective brain maps are consistent with results of other meta-analysis. The activation map of Cluster 1 (*Knowledge Representation and Language Processing*; Fig. 4a) is similar to other published meta-analytic maps and reviews of language processing and semantic representation (Binder et al. 2009; Price 2010; Price 2012; Fedorenko & Thompson-Schill 2014; Bookheimer 2002). The activation map of Cluster 3 (*Sensation, Movement and Action*; Fig 4c) is similar to other maps from studies investigating pain localization (Perini et al. 2013; Amanzio et al. 2013; Schomers & Pulvermüller 2016; Friebel et al. 2011; Vierck et al. 2013) in addition to the somatosensory co-activation network (Smith et al. 2009). Finally, the activation map of Cluster 5 (*Decision, Emotion, and Substance Use*; Fig. 4e) is also highly similar to the map of the structures involved in different aspects of emotional processing and decision-making (Bartra et al. 2013; Lindquist 2010; Etkin & Wager 2007; Buhle et al. 2014; Phan et al. 2002).

Many meta-analyses and meta-analytic tools for neuroimaging have a common (even if unstated) goal: to help homogenize our understanding of the literature and through this homogenization help define ontologies (Poldrack & Yarkoni 2016; Poldrack et al. 2011) so that we can relate brain function to cognition. However, with many tools at our disposal, there are known biases in neuroimaging (Jennings & Van Horn 2012) and the language we use can make building such ontologies difficult. With a well defined common language and homogenization of reporting results, fields such as genomics can provide a more robust assessment of the relationship between studies and the roles of particular genetic effects (Ailem et al. 2016).

Based on the analysis of term co-occurrences in the abstracts of 10,898 neuroimaging articles, we have identified a highly reliable set of dimensions and subfields that define the underlying semantic space of the neuroimaging field. Most researchers tend to stay *within* their specialized domain (by using specific key terms common to their field) and this behavior may restrict what they can conclude and how they report their findings, because they use a preferred or required terminology. In fact, Clusters 2 (*Development, Lifespan, and Disorders*) and 5 (*Decision, Emotion, and Substance Use*), as well as Components 1 and 2 show that there are language barriers between different types of clinical and experimental studies that could preclude thorough reviews of relevant literature (see examples in Supplemental Table 16).

Because such diverse terminologies and highly specialized fields could cause researchers to overlook relevant work in domains related and unrelated to their own, two recent approaches—in addition to our own—have been proposed: Papr (McGowan et al., 2017) and MAPBOT (Yuan et al., 2017). In general, our approach, Papr, and MAPBOT all aim to help users navigate literature in an easier way and to better understand the relationships between studies. Furthermore, all of these techniques use multivariate tools based on the singular value decomposition. We describe and then compare each to our approach below.

Papr was recently released to help researchers find preprints on bioRxiv that may be of interest to them. With Papr, users can move through a semantic subspace to find articles whose abstract is similar to a target abstract, as well as locate other users with similar interest^2^. Papr provides for bioRxiv some of the mechanisms (e.g., similarity and recommendation of studies) that our approach does for Neurosynth. There are, however, several major differences between our approach and Papr. First, Papr is a tool for bioRxiv while our study and many of our analyses are specifically tailored to the functional neuroimaging literature (covered by Neurosynth). Second, Papr emphasis is on study similarity. While our approach emphasizes study similarity and high-level organization of the functional neuroimaging literature, we also use the terms. The difference between Papr and our approach comes a difference of multivariate method used: Papr uses PCA whereas we use CA. CA is a bifactor technique suited to jointly accommodate the rows (studies) and columns (terms) of a matrix. Also we took the analysis of the semantic subspace further than Papr by clustering the literature into high-level domains in order to illustrate the broad configuration of the functional neuroimaging literature. Finally, while our study emphasizes the studies and words, our approach was designed around many aspects of Neurosynth, especially voxel information.

Like our study, MAPBOT utilized the Neurosynth database. MAPBOT helps researchers navigate relevant studies in Neurosynth, but conditional to a region of interest. For example, in their paper, Yuan et al., (2017) use a thalamic mask to generate a voxel × term matrix. MAPBOT extracts only the studies in Neurosynth that report voxels within an *a priori* mask to create a voxel × term matrix. MAPBOT then decomposes that voxel × term matrix with non-negative matrix factorization. MAPBOT’s goal is to provide better parcellation of regions, with richer content (i.e., terms) to help researchers understand, for examples, the functional or behavioral associations with particular parcellations within a mask. There are several major differences between our approach and MAPBOT. First is that MAPBOT analyzes voxel × term content. However, MAPBOT is restricted to *a priori* masks; that is, users must select a specific partition of voxel space. By doing so, MAPBOT cannot detect similar semantic content across voxel content. Our approach first analyzes studies × term content, and then projects (predicts) voxel content. Our approach incorporates studies, terms, and voxels for all available studies as opposed to a specific subset.

In summary: Papr is a tool to assess semantic similarity between abstracts in bioRxiv, MAPBOT parcellates *a priori* defined brain regions by using semantic content, whereas our approach assesses semantic similarity, partitions (clusters) the semantic subspace, then predicts voxel data from the semantic subspace, and finally assigns voxels to particular clusters. While both Papr and MAPBOT provide some tools to better navigate and search the literature both are lacking the key features and information we provide here. We believe that our approach to structuring the functional neuroimaging literature, and our current version of a recommendation engine, is critical to both help organize the field and to help researchers navigate the literature.

## Conclusions

To conclude, our work shows that different domains use different patterns of words, and that studies within these domains also report (or perhaps only study specific but) common brain areas. We believe that neuroimaging—and all of the domains that use and contribute to neuroimaging—would benefit from a broader harmonization of their terminology (*à la* the COBIDAS appendix on how to report routine *f*MRI analyses; Nichols et al., 2016) to put the field on the path towards formal ontologies (Poldrack & Yarkoni 2016). However, there are barriers to achieve such ontologies (see examples in Supplemental Table 16). One such barrier is time and it poses difficult questions, such as should we go back to older papers and “correct” terminology (e.g., addiction vs. substance use disorder). Another barrier is language itself because many terms have a variety of uses across disciplines (e.g., to recollect) and the same concepts could have multiple terms and used in different ways depending on factors such as stylistic choices by the authors (e.g., marijuana and cannabis). Another limitation is that some of the automated language tools commonly used (including by us) cannot always detect that certain stems have the same meaning (hippocampi vs. hippocampus). Formal and more rigorous ontologies—such as those in genomics—as well as tools more sensitive to the peculiarities of language will be required as our field moves forward and connects brain imaging to a variety of other modalities (e.g., genetics; Cioli et al. 2014; Rizzo et al. 2016), but will require effort from a variety of disciplines to harmonize and standardize terminology.

## Acknowledgements

We would like to thank Micaela Chan and Jenny Rieck for their feedback on previous versions of this manuscript. We also greatly appreciate all the public commentary we received provided by many individuals in various formats (e.g., Twitter).

## Footnotes

1 We used absolute correlation because there can be trivial sign flips between sub-samples of data, so the sign is irrelevant but the magnitude of the correlation is relevant.

2 Currently there is an offline version of Papr here: https://github.com/jtleek/papr. During the writing of our manuscript a “live” version of Papr was available but is no longer: https://jhubiostatistics.shinyapps.io/papr/.

